# FAIRSCAPE: A Framework for FAIR and Reproducible Biomedical Analytics

**DOI:** 10.1101/2020.08.10.244947

**Authors:** Maxwell Adam Levinson, Justin Niestroy, Sadnan Al Manir, Karen Fairchild, Douglas E. Lake, J. Randall Moorman, Timothy Clark

## Abstract

Results of computational analyses require transparent disclosure of their supporting resources, while the analyses themselves often can be very large scale and involve multiple processing steps separated in time. Evidence for the correctness of any analysis should include not only a textual description, but also a formal record of the computations which produced the result, including accessible data and software with runtime parameters, environment, and personnel involved.

This article describes FAIRSCAPE, a reusable computational framework, enabling simplified access to modern scalable cloud-based components. FAIRSCAPE fully implements the FAIR data principles and extends them to provide fully FAIR Evidence, including machine-interpretable provenance of datasets, software and computations, as metadata for all computed results.

The FAIRSCAPE microservices framework creates a complete Evidence Graph for every computational result, including persistent identifiers with metadata, resolvable to the software, computations, and datasets used in the computation; and stores a URI to the root of the graph in the result’s metadata. An ontology for Evidence Graphs, EVI (https://w3id.org/EVI), supports inferential reasoning over the evidence.

FAIRSCAPE can run nested or disjoint workflows and preserves provenance across them. It can run Apache Spark jobs, scripts, workflows, or user-supplied containers. All objects are assigned persistent IDs, including software. All results are annotated with FAIR metadata using the evidence graph model for access, validation, reproducibility, and re-use of archived data and software.

## Introduction

### 1. Motivation

Computation is an integral part of the preparation and content of modern biomedical scientific publications, and the findings they report. Computations can range in scale from simple statistical routines run in Excel spreadsheets to massive orchestrations of very large primary datasets, computational workflows, software, cloud environments, and services. They typically produce data and generate images or tables as output. Scientific claims of the authors are supported by evidence that includes reference to the theoretical constructs embodied in existing domain literature, and to the experimental or observational data and its analysis represented in images or tables.

Today, increasingly strict requirements are demanded to leave a digital footprint of each preparation and analysis step in derivation of a finding to support reproducibility and reuse of both data and tools. The widely recommended and often required practice by publishers and funders today is to archive and cite one’s own experimental data (Cousijn et al. 2018; Data Citation Synthesis Group 2014; Fenner et al. 2019; Groth et al. 2020); and to make it FAIR (Wilkinson et al. 2016). These approaches were developed over more than a decade by a significant community of researchers, archivists, funders, and publishers, prior to the current recommendations (Altman et al. 2001; Altman and King 2007; Borgman 2012; Bourne et al. 2012; Brase 2009; CODATA/ITSCI Task Force on Data Citation 2013; King 2007; Starr et al. 2015; Uhlir 2012). There is increasing support among publishers and the data science community to recommend, in addition, archiving and citing the specific software versions used in analysis (D. Katz et al. 2021; Smith et al. 2016), with persistent identification and standardized core metadata, to establish FAIRness for research software (D. S. Katz et al. 2021; Lamprecht et al. 2020); and to require identification via persistent identifiers, of critical research reagents (A. Bandrowski 2014; A. E. Bandrowski and Martone 2016; Prager et al. 2018).

How do we facilitate and unify these developments? Can we make the recorded digital footprints as broadly useful as possible in the research ecosystem, while their generation occurs as side-effects of processes inherently useful to the researcher – for example, in large scale data analytics and data commons environments?

The solution we developed is a *reusable framework for building provenance-aware data commons environments*, which we call FAIRSCAPE. It provides several features directly useful to the computational scientist, by simplifying and accelerating important data management and computational tasks; while providing, as metadata, an integrated **evidence graph** of the resources used in performing the work, allowing them to be retrieved, validated, reused, modified, and extended.

Evidence graphs are formal models inspired by a large body of work in abstract argumentation (Bench-Capon and Dunne 2007; Brewka et al. 2014; Carrera and Iglesias 2015; Cayrol and Lagasquie-Schiex 2009; Dung 1995; Dung and Thang 2018; Gottifredi et al. 2018; Rahwan 2009; Schneider et al.), and analysis of evidence chains in biomedical publications (Tim Clark et al. 2014; Greenberg 2009, 2011), which shows that the evidence for correctness of any finding, can be represented as a directed acyclic support graph, an Evidence Graph. When combined with a graph of challenges to statements, or their evidence, this becomes a bipolar argument graph - or argumentation system (Cayrol and Lagasquie-Schiex 2009, 2010, 2013).

The nodes in these graphs can readily provide metadata about the objects related to the computation, including the computation parameters and history. Each set of metadata may be indexed by one or more persistent identifiers, as specified in the FAIR principles; and may include a URI by which the objects themselves may be retrieved, given the appropriate permissions. In this model, core metadata retrieved on resolution of a persistent identifier (PID) (Juty et al. 2020; Starr et al. 2015) will include an evidence graph for the object referenced by the PID. A link to the object’s evidence graph can be embedded in its metadata.

The central goals of FAIRSCAPE can be summarized as (1) to develop reusable cloud-based “data commons” frameworks adapted for very large-scale data analysis, providing significant value to researchers; and (2) to make the computations, data, and software in these environments fully transparent and FAIR (findable, accessible, interoperable, reusable). FAIRSCAPE supports a “data ecosystem” model (Grossman 2019) in which computational results and their provenance are transparent, verifiable, citable, and FAIR across the research lifecycle. We combined elements of prior work by ourselves and others on provenance, abstract argumentation frameworks, data commons models, and citable research objects, to create the FAIRSCAPE framework. This work very significantly extends and refactors the identifier and Metadata Services we and our colleagues developed in the NIH Data Commons Pilot Project Consortium (Timothy Clark et al. 2018; Fenner et al. 2018; *NIH Data Commons Pilot: Object Registration Service (ORS)* 2018)

FAIRSCAPE has a unique position in comparison to other provenance-related, reproducibility-enabling, and “data commons” projects. We combine elements of all three approaches, while providing transparency, FAIRness, validation, and re-use of resources; and emphasize reusability of the FAIRSCAPE platform itself. Our goal is to enable researchers to implement effective and useful provenance-aware computational data commons in their own research environments, at any scale, while supporting full transparency of results across projects, via Evidence Graphs represented using a formal ontology.

### 2. Related Work

Works focusing on provenance per se such as (Alterovitz et al. 2018; Ellison et al. 2020) and the various workflow provenance systems such as (Khan et al. 2019; Papadimitriou et al. 2021; Yakutovich et al. 2021) are primarily concerned with very detailed documentation of each computation on one or more datasets. The W3C PROV model (Gil et al. 2013; Lebo et al. 2013; Moreau et al. 2013) was developed initially to support interoperability across the transformation logs of workflow systems. Our prior work on Micropublications (Tim Clark et al. 2014) extending and repurposing several core classes and predicates from W3C PROV, were preliminary work forming a basis for the EVI ontology (Al Manir et al. 2021a, 2021b).

The EVI ontology used in FAIRSCAPE to represent evidence graphs, is concerned with creating reasonable transparency of evidence supporting scientific claims, including computational results; it reuses the three major PROV classes *Entity, Activity*, and *Agent* as a basis to develop a detailed ontology and rule system for reasoning across the evidence for (and against) results. When a computational result is reused in any new computation, that information is added to the graph, whether or not the operations were controlled by a workflow manager. Challenges to results, datasets, or methods, may also be added to the graph. While our current use of EVI is on computational evidence, it is designed to be extensible to objects across the full experimental and publication lifecycle.

Systems providing data commons environments, such as the various NCI and NHLBI cloud platforms (Birger et al. 2017; Brody et al. 2017; Lau et al. 2017; Malhotra et al. 2017; Wilson et al. 2017) while providing many highly useful specialized capabilities for their domain users, including re-use of data and software, have not focused extensively on providing re-use of their own frameworks, and are centralized. As noted later in this article, FAIRSCAPE can be – and is meant to be - installed on public, private, or hybrid cloud platforms, “bare metal” clusters, and even on high-end laptops, for use at varying scopes – personal, institution-wide, lab-wide, multi-center, etc.

Reproducibility platforms such as Whole Tale and CodeOcean, (Brinckman et al. 2019; Chard et al. 2019; Merkys et al. 2017) attempt to take on a one-stop-shop role for researchers wishing to demonstrate or at least assert, reproducibility of their computational research. Of these, CodeOcean (https://codeocean.com) is a special case – it is run by a company and appears to be principally described in press releases, and not in any peer reviewed articles.

FAIRSCAPE’s primary goals are to enable construction of multi-scale computational data lakes, or commons; and to make results transparent for reuse across the digital research ecosystem, via FAIRness of data, software, and computational records. FAIRSCAPE supports reproducibility via transparency.

In very many cases - such as the very large analytic workflows in our first use case - we believe that no reviewer will attempt to replicate such large-scale computations, which ran for months on substantial resources. The primary use case will be validation via inspection, and *en passant* validation via software reuse.

FAIRSCAPE is not meant to be a one-stop shop. It is a transferable, reusable framework. It is not only intended to enable localized participation in a global, fully FAIR data and software ecosystem – it is itself FAIR software. The FAIRSCAPE software, including installation and deployment instructions, is available in the CERN Zenodo archive (Levinson et al. 2021); and in the FAIRSCAPE Github repository (https://github.com/fairscape/fairscape).

#### Enabling Transparency through EVI’s Formal Model

To enable the necessary results transparency across separate computations, we abstracted core elements of our micropublications model (Tim Clark et al. 2014) to create EVI (http://w3id.org/EVI), an ontology of evidence relationships that extends W3C PROV to support specific evidence types found in biomedical publications; and enable reasoning across deep evidence graphs, and propagation of evidence challenges deep in the graph, such as: retractions, reagent contamination, errors detected in algorithms, disputed validity of methods, challenges to validity of animal models, and others. EVI is based on the fundamental idea that scientific findings or claims are not facts, but assertions backed by some level of evidence, *i.e*., they are defeasible components of argumentation. Therefore, EVI focuses on the structure of evidence chains that support or challenge a result, and on providing access to the resources identified in those chains. Evidence in a scientific article is in essence, a record of the provenance of the finding, result, or claim asserted as likely to be true; along with the theoretical background material supporting the result’s interpretation.

If the data and software used in analysis are all registered and receive persistent identifiers (PIDs) with appropriate metadata, a provenance-aware computational data lake, *i.e*., a data lake with provenance-tracking computational services, can be built that attaches evidence graphs to the output of each process. At some point, a citable object - a dataset, image, figure, or table will be produced as part of the research. If this, too, is archived with its evidence graph as part of the metadata and the final supporting object is either directly cited in the text, or in a figure caption, then the complete evidence graph may be retrieved as a validation of the object’s derivation and as a set of URIs resolvable to reusable versions of the toolsets and data. Evidence graphs are themselves entities that can be consumed and extended at each transformation or computation.

The remainder of this article describes the approach, microservices architecture, and interaction model of the FAIRSCAPE framework in detail.

## Materials and Methods

### 1. FAIRSCAPE Architectural Layers

FAIRSCAPE is built on a multi-layer set of components using a containerized microservice architecture (MSA) (Balalaie et al. 2016; Larrucea et al. 2018; Lewis and Fowler 2014; Wan et al. 2018) running under Kubernetes (Burns et al. 2016). We run our local instance in an OpenStack (Adkins 2016) private cloud environment, and maintain it using a DevOps deployment process (Balalaie et al. 2016; Leite et al. 2020). FAIRSCAPE may also be installed on laptops running minikube in Ubuntu Linux, MacOS, or Windows environments; and on Google Cloud managed Kubernetes. An architectural sketch of this model is shown in Figure 1.

**Figure 1.**
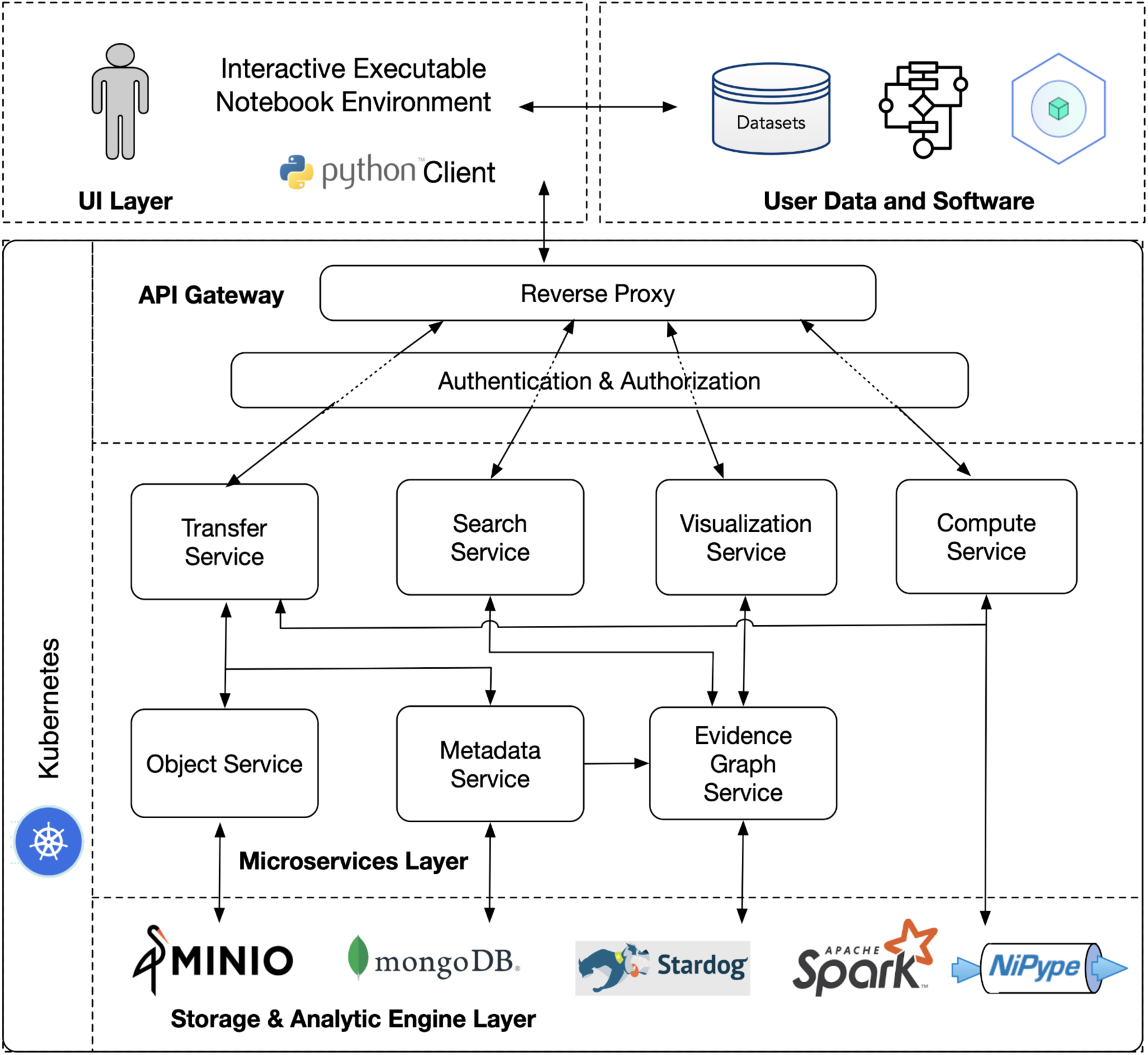
FAIRSCAPE architectural layers and components.

Ingress to microservices in the various layers is through a reverse proxy using an API gateway pattern. The top layer provides an interface to the end users with raw data and the associated metadata. The mid layer is a collection of tightly coupled services that allow end users with proper authorization to submit and view their data, metadata, and various types of computations performed on them. The bottom layer is built with special purpose storage and analytics platforms for storing and analyzing data, metadata and provenance information. All objects are assigned PIDs using local ARK (Kunze and Rodgers 2008) assignment for speed, with global resolution for generality.

#### 1.1 UI Layer

The User Interface layer in FAIRSCAPE offers end users various ways to utilize the functionalities in the framework. A Python client simplifies calls to the microservices. Data, metadata, software, scripts, workflows, containers, etc. are all submitted and registered by the end users from the UI Layer, which may be configured to include an interactive executable notebook environment such as Binder or Deepnote.

#### 1.2 API Gateway

Access to the FAIRSCAPE environment is through an API gateway, mediated by a reverse proxy. Our gateway is mediated by Traefik (https://traefik.io) which dispatches calls to the various microservices endpoints.

Traefik is a reverse proxy that we configure as a Kubernetes *Ingress Controller*, to dynamically configure and expose multiple microservices using a single API.

The endpoints of the services are exposed through the OpenAPI specification (formerly Swagger Specification) (Darrel Miller et al. 2020) which defines the standard and the language-agnostic interface for publishing RESTful APIs and allows service discovery. Accessing the services requires user authentication, which we implement using the Globus Auth authentication broker (Tuecke et al. 2016). Users of GlobusAuth may be authenticated via a number of permitted authentication services, and are issued a token which serves as an identity credential. In our current installation we require use of the CommonShare authenticator, with site-specific two-factor authentication necessary to obtain an identity token. This token is then used by the microservices to determine a user’s permission to access various functionality.

#### 1.3 Authentication and Authorization Layer

Authentication and authorization (authN/authZ) in FAIRSCAPE are handled by Keycloak (Christie et al. 2020), a widely-used open source identity and access management tool.

When Traefik receives a request, it handles an authentication check to Keycloak, which then determines whether or not the requestor has a valid token for an identity that can perform the requested action.

We distribute FAIRSCAPE with a preconfigured Keycloak for basic username / password authentication & authorization of service requests. This can be easily modified to support alternative identity providers, including LDAP, OpenID Connect, and OAuth2.0 for institutional single sign-on. Services continue to interact the same way, even if you change the configured identity provider.

Within our local Keycloak configuration, we chose to define Globus Auth as the identity provider. Globus Auth then serves as a dispatching broker amongst multiple other possible final identity providers. We selected the login service at the University of Virginia as our final provider, providing two-factor authentication and institutional single sign-on. Keycloak is very flexible in allowing selection of various authentication schemes, such as LDAP, SAML, OAuth2.0, etc. Selection of authentication schemes is an administrator decision.

#### 1.4 Microservices Layer

The microservices layer is composed of seven services: (1) Transfer, (2) Metadata, (3) Object, (4) Evidence Graph, (5) Compute, (6) Search, and (7) Visualization. These are described in more detail in Section 2. Each microservice does its own request authorization, subsequent to Keycloak, enabling fine-grained access control.

#### 1.5 Storage and Analytic Engine Layer

In FAIRSCAPE, an S3 compatible object store is required for storing objects, a document store for storing metadata, and a graph store for storing graph data. Persistence for these databases is configured through Kubernetes volumes, which map specific paths on containers to disk storage. The current release of FAIRSCAPE uses the S3 compatible MinIO as the object store, MongoDB as the document store, and Stardog as the graph store. Computations invoked by the Compute Service are managed by Kubernetes, Apache SPARK, and the Nipype neuroinformatics workflow engine.

### 2. FAIRSCAPE Microservice Components

#### 2.1 Transfer Service

This service transfers and registers digital research objects - datasets, software, etc., - and their associated metadata, to the Commons. These objects are sent to the transfer service as binary data streams, which are then stored in MinIO object storage. These objects may include structured or unstructured data, application software, workflow, scripts. The associated metadata contains essential descriptive information such as context, type, name, textual description, author, location, checksum, etc. about these objects. Metadata are expressed as JSON-LD and sent to the Metadata Service for further processing.

Hashing is used to verify correct transmission of the object – users are required to specify a hash which is then recomputed by the Object Service after the object is stored. Hash computation is currently based on the SHA-256 secure cryptographic hash algorithm (Dang 2015). Upon successful execution, the service returns a PID of the object in the form of an ARK, which resolves to the metadata. The metadata includes, as is normal in PID architecture (Starr et al. 2015), a link to the actual data location.

An OpenAPI description of the interface is here: https://app.swaggerhub.com/apis/FAIRSCAPE/Transfer/0.1

#### 2.2 Metadata Service

The Metadata Service handles metadata registration and resolution including identifier minting in association with the object metadata. The Metadata Service takes user POSTed JSON-LD metadata and uploads the metadata to MongoDB and Stardog, and returns a PID. To retrieve metadata for an existing PID a user makes a GET call to the service. A PUT call to the service will update an existing PID with new metadata. While other services may read from MongoDB and Stardog directly, the Metadata Service handles all writes to MongoDB and Stardog.

An OpenAPI description of the interface is here: https://app.swaggerhub.com/apis/FAIRSCAPE/Metadata-Service/0.1

#### 2.3 Object Service

*The Object Service provides a direct interface between the Transfer Service and MinIO as well as maintaining consistency between MinIO and the metadata store. The Object Service handles uploads of new objects as well as uploading new versions of existing files. In both cases the Object Service accepts a file and desired file location as inputs and (if the location is available) uploads the file to desired location in MinIO and returns a PID representing the location of the uploaded file. A DELETE call to the service will delete the requested file from MinIO as well as delete the PID with the link to the data, however the PID representing the object metadata remains*.

An OpenAPI description of the interface is here: *https://app.swaggerhub.com/apis/FAIRSCAPE/Object-Service/0.1*

#### 2.4 Evidence Graph Service

The Evidence Graph Service creates a JSON-LD Evidence Graph of all provenance related metadata to a PID of interest. The Evidence Graph documents all objects such as datasets, software, workflows, and the computations which are directly involved in creating the requested entity. The service accepts a PID as its input, runs a PATH query built on top of the SPARQL query engine in Stardog with the PID of interest as its source to retrieve all supporting nodes. To retrieve an Evidence Graph for a PID a user may make a GET call to the service.

The Evidence Graph Service plays an important role in reproducing computations. All resources required to run a computation are exposed using persistent identifiers by the evidence graph. A user can reproduce the same computation by invoking the appropriate services available through the Python client with the help of these identifiers. This feature allows a user to verify the accuracy of the results and detect any discrepancies.

An OpenAPI description of the interface is here: https://app.swaggerhub.com/apis/FAIRSCAPE/Evidence-Graph/0.1

#### 2.5 Compute Service

This service executes user uploaded scripts, workflows, or containers, on uploaded data. It currently offers two compute engines (Spark, Nipype) in addition to native Kubernetes container execution, to meet a variety of computational needs. Users may execute any script they would like to run as long as they provide a docker container with the required dependencies. To complete jobs the service spawns specialized pods on Kubernetes designed to perform domain specific computations that can be scaled to the size of the cluster. This service provides the essential ability to recreate computations based solely on identifiers. For data to be computed on it must first be uploaded via the Transfer Service and be issued an associated PID.

The service accepts a PID for a dataset, a script, software, or a container, as input and produces a PID representing the activity to be completed. The request returns a job identifier from which job progress can be followed. Upon completion of a job all outputs are automatically uploaded and assigned new PIDs, with provenance aware metadata. At job termination, the service performs a ‘cleanup’ operation, where a job is removed from the queue once it is completed.

An OpenAPI description of the interface is here: https://app.swaggerhub.com/apis/FAIRSCAPE/Compute/0.1

#### 2.6 Search Service

The Search Service allows users to search for object metadata containing strings of interest. It accepts a string as input and performs a search over all literals in the metadata for exact string matches and returns a list of all PIDs with a literal containing the query string. It is invoked via the GET method of API endpoint to the service with the search string as argument.

An OpenAPI description of the interface is here: https://app.swaggerhub.com/apis/FAIRSCAPE/Search/0.1

#### 2.7 Visualization Service

This service allows users to visualize Evidence Graphs interactively in the form of nodes and directed edges, offering a consolidated view of the entities and the activities supporting correctness of the computed result. Our current visualization engine is Cytoscape (Shannon 2003). Each node displays its relevant metadata information, including its type and PID, resolved in real-time.

The Visualization Service renders the graph on an HTML page.

An OpenAPI description of the interface is here: https://app.swaggerhub.com/apis/FAIRSCAPE/Visualization/0.1

### 3. FAIRSCAPE Service Orchestration

FAIRSCAPE orchestrates a set of containers to provide patterns for object registration, including identifier minting and resolution; object retrieval; computation; search; evidence graph visualization, and object deletion. These patterns are orchestrated following API ingress, authentication, and service dispatch, by microservice calls, invoking the relevant service containers.

#### 3.1 Object Registration

Object registration occurs initially via the Transfer Service, with an explicit user service call, and again automatically using the same service, each time a computation generates output. Objects in FAIRSCAPE may be software, containers, or datasets. Descriptive metadata must be specified for object registration to occur.

When invoked, the Transfer Service calls the Metadata Service (MDS) to mint a new persistent identifier, implemented as an Archival Resource Key (ARK), generated locally, and to store it associated with the descriptive metadata, including the new registered object location. MDS stores object metadata, including provenance, in both MongoDB and in the Stardog graph store, allowing subsequent access to the object metadata by other services.

After minting an identifier and storing the metadata, the Transfer Service calls the Object Service to persist the new object, and then updates the metadata with the stored object location. Hashing is used to verify correct transmission of the object – users are required to specify a SHA256 hash on registration, which is then recomputed by the Object Service and verified after the object is stored. Internally computed hashes are provided for re-verification when the object is accessed. Failure of hashes to match generates an error.

##### 3.1.1 Identifier Minting

The Metadata Service mints PIDs in the form of ARKs. Multiple alternative PIDs may exist for an object and PIDs are resolved to their associated object level metadata including the object’s Evidence Graph and location with appropriate permissions.

In the current deployment, ARKs created locally are registered to an assigned Name Assigning Authority Number. The ARK globally unique identifier ecosystem employs a flexible minimalistic standard and existing infrastructure.

##### 3.1.2 Identifier Resolution

ARK identifier resolution may be handled locally and/or by external resolver services such as Name-to-Thing (https://n2t.net). The Name-to-Thing resolver allows for Name Assigning Authority Numbers (NAAN) to have redirect rules for their ARKs, which forwards requests to the Name Mapping Authority Hostport for the corresponding commons. Each FAIRSCAPE instance should independently obtain a NAAN, and a DNS name for their local FAIRSCAPE installation, if they wish their ARKs to be resolved by n2t.net. DataCite DOI registration and resolution are planned for future work.

#### 3.2 Object Retrieval

Objects are accessed by their PID, after prior resolution of the object’s PID to its metadata (MDS) and authorization of the user’s authentication token for data access on that object. Object access is either directly from the Object Store, or from wherever else the object may reside. Certain large objects residing in robust external archives, may not be acquired into local object storage, but remain in place, up to the point of computation.

#### 3.3 Computation

When executing a workload through the compute service, data, software, and containers are referenced through their PIDs, and by no other means. The Compute Service utilizes the stored metadata to dereference the object locations, and transfers them to the managed containers. The compute service also creates a provenance record of its own execution, associated with an identifier of type evi:Computation. Upon the completion of a job, the Compute Service stores the generated output through the Transfer Service. Running workloads in the Compute Service enables all data, results, and methods to be tracked via a connected evidence graph, with persistent identifiers available for every node.

The Compute Service executes computations using (a) a container specified by the user, or (b) the Apache Spark service, or (c) the Nipype workflow engine. Like datasets and software (including scripts), computations are represented by persistent identifiers assigned to them. Objects are passed to the Compute Service by their PIDs and the computation is formally linked to the software (or script) by the usedSoftware property, and the input datasets by the usedDataset property.

Runtime parameters may be passed with objects and a single identifier is minted with the given parameters and connected to the computation via the ‘parameters’ property. However, at this moment these parameters are not incorporated in the evidence graph.

The Compute Service spawns a Kubernetes pod with the input objects mounted in the /data directory by default. Upon completion of the job all output files in the /outputs directory are transferred to the object store and identifiers for them are minted with the property generatedBy. The generatedBy property references the identifier for the computation.

#### 3.4 Object Search

Object searches are performed by the Search Service, called directly on Service Dispatch. Search makes use of Stardog’s full text retrieval, which in turn is based on Apache Lucene.

#### 3.5 Evidence Graph Visualization

Evidence graphs of any object acquired by the system may be visualized at any point in this workflow using the Visualization Service. Nipype provides a chart of the workflows it executes using the Graphviz package. Our Evidence Graph Service is interactive, using the Cytoscape package (Shannon 2003), and allows Evidence graphs of multiple workflows in sequence to be displayed whether or not they have been combined into a single flow.

#### 3.6 Object Deletion

Objects are deleted by calls to the Object Service to clear the object from storage, which then calls MDS and nulls out the object location in the metadata record. Metadata is retained even though the object may cease to be held in the system, in accordance with the Data Citation Principles (Data Citation Synthesis Group 2014).

## Results

Two use cases are presented here to demonstrate the application of FAIRSCAPE services. The first use case performs analysis of time series algorithms while the second runs a neuroimaging workflow. For each use case, the operations involving data transfer, computation, and evidence graph generation are described below:

### 1. Use Case Demonstration 1: Highly Comparative Time Series Analysis (HCTSA) of NICU Data

Researchers at the Neonatal Intensive Care Unit (NICU) at the University of Virginia continuously monitor infants and collect vital signs such as heart rate (HR) and oxygen saturation. Patterns in vital sign measurements may indicate acute or chronic pathology among infants. In the past a few standard statistics and algorithms specially designed for discovering certain pathologies were applied to similar vital sign data. In this work we additionally applied many time series algorithms from other domains with the hope that these algorithms would be helpful for prediction of unstudied outcomes. A total of 67 time series algorithms have been recoded as Python scripts and run on the vital signs of 5,997 infants collected over 10 years during 2009-2019. The data are then merged, sampled and clustered to find representative time series algorithms which express unique characteristics about the time series and make it easier for researchers to quickly build models for outcomes where the physiology is not known. FAIRSCAPE can be used to build such models using its services.

A series of steps are required to execute and reproduce the HCTSA of NICU data. They include transferring NICU data, Python scripts, and associated metadata to the storage to be used later, running the scripts in the FAIRSCAPE compute environment, and generating the evidence graph. The first script runs the time series algorithms on the vital sign data while the second script performs clustering of these algorithms to generate a heatmap image. The evidence graph generated for all patients contains over 17,000 nodes. However, a simplified version of the computational analysis based on a single patient is described here as the steps for executing the analysis are common for all patients. These steps are briefly described below:

#### 1.1 Transfer Data, Software and Metadata

Before any computation is performed FAIRSCAPE requires each piece of data and software to be uploaded to the object store with its metadata using the POST method of the transfer service. The raw data file containing the vital signs and the associated metadata are uploaded first. The scripts and the associated metadata are uploaded next.

The `upload_file` function shown below is used to transfer the raw data file `UVA_7129_HR.csv` as the first parameter and the associated metadata referenced by the variable `dataset_meta` as the second parameter:

~~~
dataset_meta = {
 “@context”:{
 “@vocab”:”http://schema.org/“
 },
 “@type”:”Dataset”,
 “name”:”Raw Data”,
 “description”:”Heart Rate Measures from patient 7129 from admission to discharge.”
}
raw_ts_fs_id = FAIR.upload_file(‘UVA_7129_HR.csv’, dataset_meta)
~~~

As part of the transfer, identifiers are minted by the Metadata Service for each successfully uploaded object. The variable `raw_time_series_fs_id` refers to the minted identifier returned by the function. Each identifier is resolvable to the uploaded object which can be accessed only by an authorized user.

#### 1.2 Time Series Data Analysis

Once the transfer is complete, the computation for the data analysis can be started. The computation takes the raw vital sign measurements as input, groups the measurements into 10-minute intervals and runs each algorithm on them. FAIRSCAPE makes launching such a computation easy by executing the POST method of the compute service with identifiers of the data and script as parameters. The compute service creates an identifier with metadata pointing to the provided inputs and launches a Kubernetes pod to perform the computation. Upon completion of the script, all output files are assigned identifiers and stored in the object store.

The `compute` function takes the PIDs of the dataset, software/script, and the type of job such Apache Spark, Nipype, or custom containers as parameters. The PID it returns refers to the submitted job which can be used to track the progress of the computation and its outputs.

The `compute` function shown below is used to launch a computation on the raw data file `raw_ts_fs_id` as the first parameter, the analysis script `raw_data_analysis_script_id` as the second parameter using their identifiers, and the type of job `spark` as the third parameter:

~~~
raw_data_analysis_job_id = FAIR.compute(raw_ts_fs_id, raw_data_analysis_script_id, ‘spark’)
~~~

The PID the compute function returns resolves to the submitted job and is referenced by

~~~
raw_data_analysis_job_id.
~~~

#### 1.3. Clustering of Algorithms

The next computation step is to perform clustering of algorithms. Many algorithms are from similar domains and the operations they perform express similar characteristics. The HCTSA Clustering script clusters these algorithms into groups which are highly correlated and a representative algorithm could be chosen from each grouping. The compute service is then invoked with identifiers of the clustering script and the processed data as parameters. The compute function below takes PIDs of the processed time series feature set, clustering script, and spark job type as input parameters and returns a PID representing the job to generate the HCTSA heatmap:

~~~
HCTSA_Heatmap_job_id = FAIR.compute(processed_ts_fs_id, hctsa_clustering_script_id, ‘spark’)
~~~

An image showing the clustered algorithms is produced at the end of this step which is shown in Figure 2.

**Figure 2.**
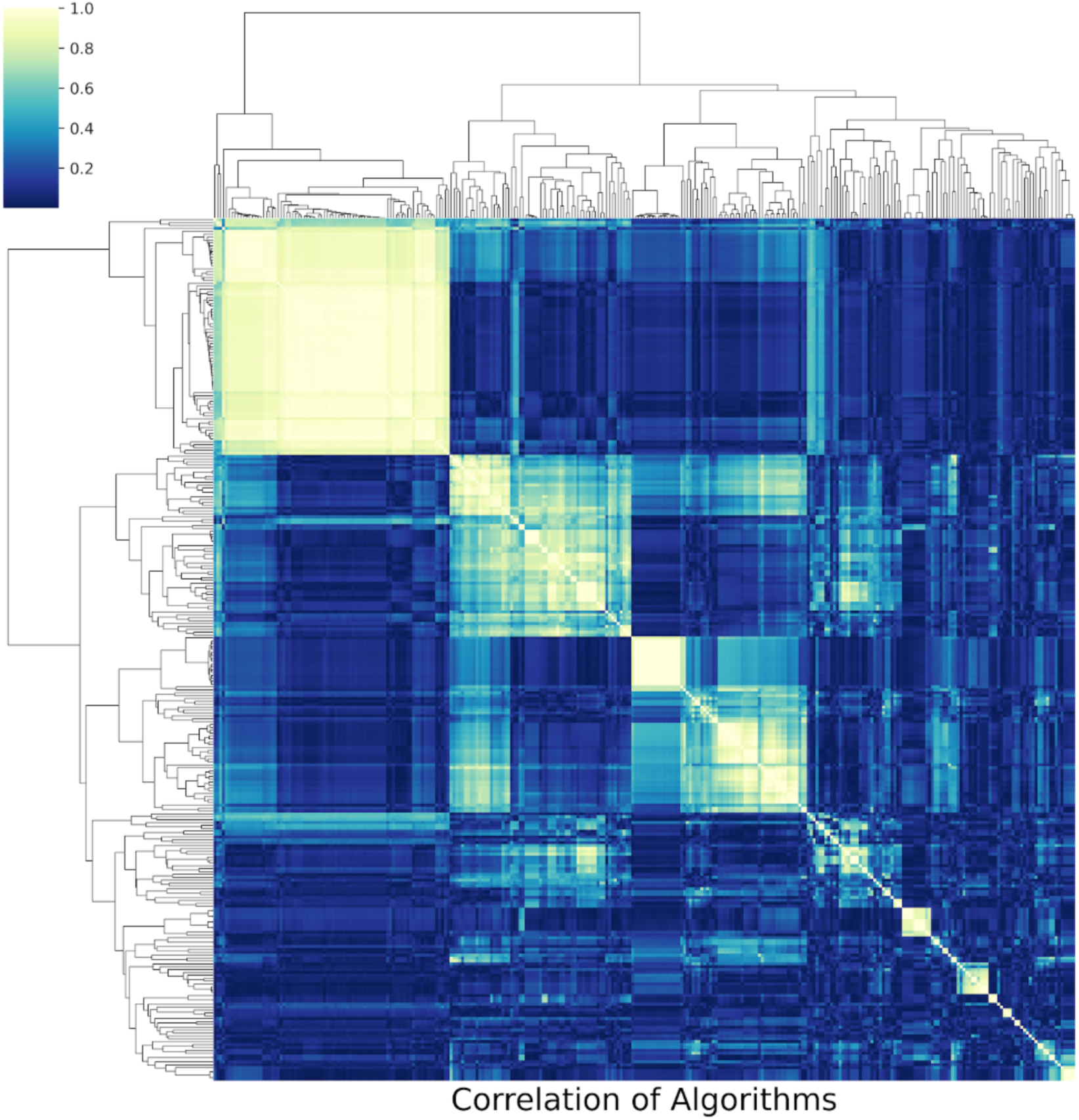
NICU HCTSA clustering heatmap. *X axis and Y axis are operations (algorithms using specific parameter sets), color is correlation between algorithms. The large white squares are clusters of highly correlated operations which suggest the dimension of the data may be greatly diminished by selecting “representative” algorithms from these clusters*.

#### 1.4. Generating the Evidence Graph

An evidence graph is generated using the GET method of the Evidence Graph Service. Figure 3 illustrates all computations and the associated inputs and outputs for a single patient. The graph for all patients contains 17,995 nodes of types *Image, Computation, Dataset* and *Software*. Each patient has a unique Raw Time Series Feature Set, a Raw Data Analysis computation, and a Processed Time Series file. The Raw Data Analysis Script, HCTSA Clustering Script, HCTSA Heatmap Generation, and HCTSA Heatmap are shared among all patients.

**Figure 3.**
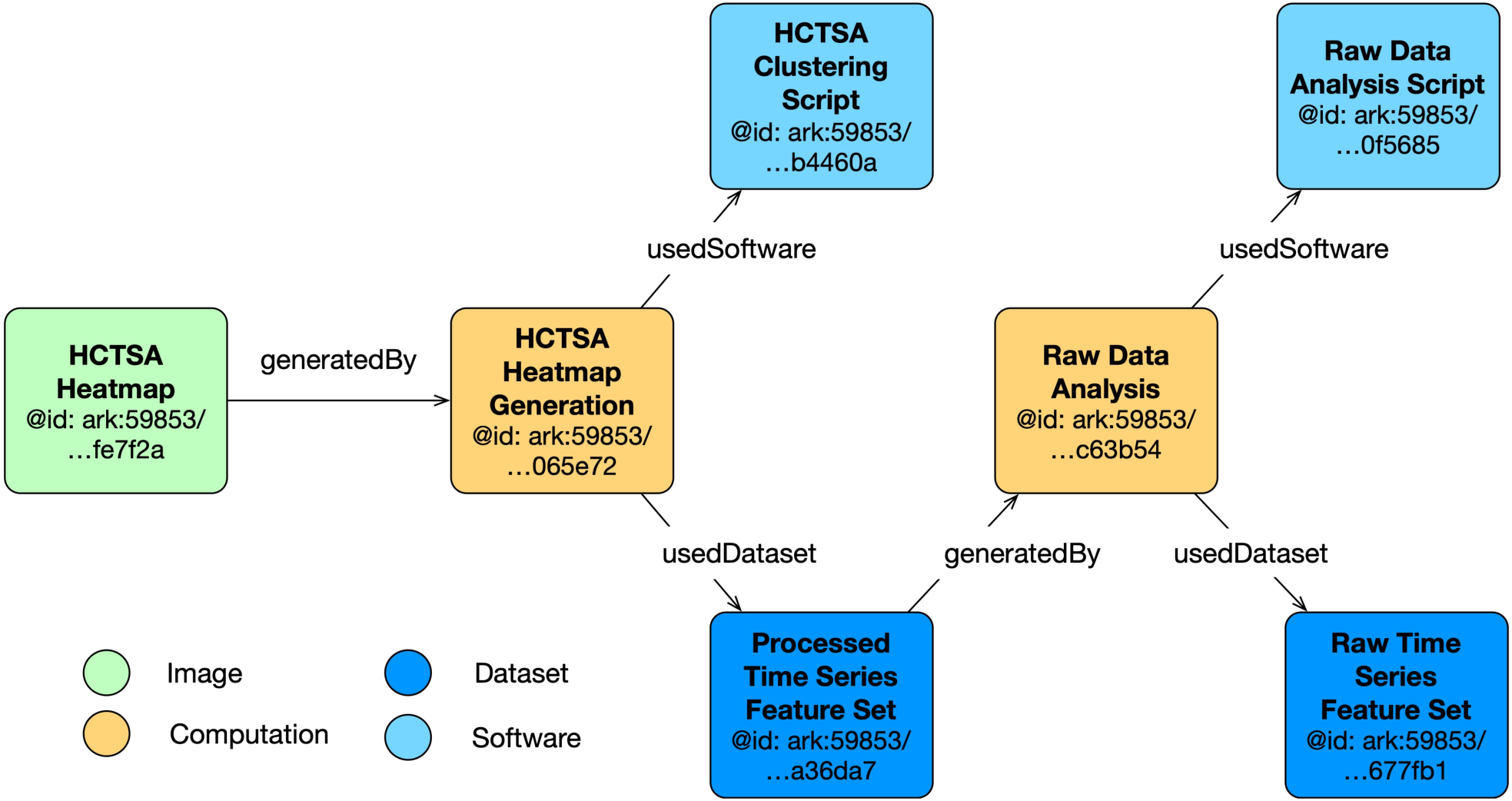
Simplified Evidence graph for one patient’s computations. *Vital signs = dark blue box bottom right; computations = yellow boxes; processed data = dark blue box in middle; green box = heatmap of correlations*.

The simplified evidence graph in Figure 3 contains 7 nodes, each with its own PID, where a Computation (Raw Data Analysis) uses a Dataset (Raw Time Series Feature Set) as the input to a Software (Raw Data Analysis Script), representing the script to execute all the time series algorithms, and generates the Dataset (Processed Time Series Feature Set) as the output. The next Computation (HCTSA Cluster Heatmap Generation) uses the processed Dataset generated during the previous computation as the input of the Software (HCTSA Clustering Script), which generates an Image (HCTSA Cluster Heatmap), representing the clustering of the algorithms as an output. The `evidence_graph` function takes the PID of the HCTSA Heatmap image:

`evidence_graph_jsonld = FAIR.evidence_graph(HCTSA_Heatmap_id)`

and generates the evidence graph for that PID serialized in JSON-LD (shown in Figure 4).

**Figure 4.**
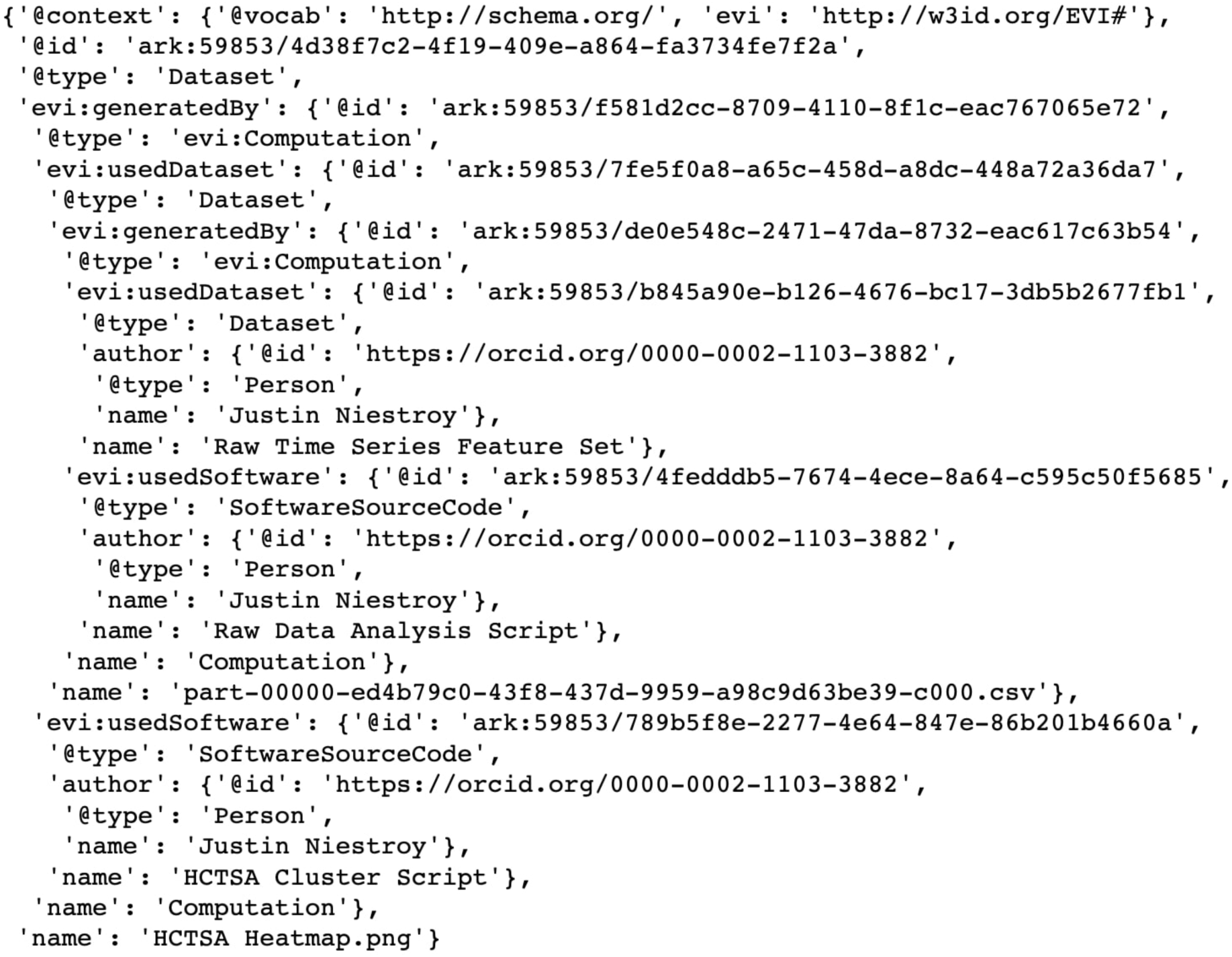
JSON-LD Evidence Graph for patient computation as illustrated in Figure 3.

### 2. Use Case Demonstration 2: Neuroimaging Analysis Using Nipype Workflow Engine

Data analysis in neuroimaging often requires multiple heterogeneous algorithms which sometimes lack transparent interoperability under a uniform platform. Workflows offer solutions to this problem by bringing the algorithms and software under a single umbrella. The open-source neuroimaging workflow engine *Nipype* combines heterogenous neuroimaging analysis software packages under a uniform operating platform which resolves the interoperability issues by allowing them to talk to each other. Nipype provides access to a detailed representation of the complete execution of a workflow consisting of inputs, output and runtime parameters. The containerization-friendly release, detailed workflow representation, and minimal effort required to modify existing services to produce a deep evidence graph have made Nipype an attractive target for integration within the FAIRSCAPE framework.

Among the services in FAIRSCAPE, only the compute service needed to be modified to run and interrogate Nipype. The modifications include repurposing the service to run the workflow from the Nipype-specific container generated by the Neurodocker tool and to capture all entities from the internal graph generated after the workflow is executed. Whereas an evidence graph typically includes the primary inputs and outputs, the deep evidence graph produced here additionally contains intermediate inputs and outputs. It provides a detailed understanding of each analysis performed using the computations, software and datasets.

A workflow is considered simple if it consists of a sequence of processing steps and complex if there is nesting of workflow execution such that the output of one workflow is used as the input to another workflow. The simple neuroimaging preprocessing workflow demonstrated here (Notter 2020) involves steps to correct motion of functional images, co-register functional images to anatomical images, smooth the co-registered functional images, and detect artifacts in functional images. As part of the data transfer, dataset containing images and the script to run the processing steps with their associated metadata were uploaded using the `file_upload` function as shown in the previous use case. The compute function is then used to launch a computation on the image dataset which runs the processing script on the repurposed Nipype container. The only exception in the `compute` function is that it uses `nipype` as the third input parameter instead of `spark` when the Compute Service was invoked. The full evidence graph, generated using the `evidence_graph` function as demonstrated above, is too large to document due to space constraints. Therefore, only the graph of the motion correction of functional images with FSL’s MCFLIRT is shown in Figure 5. For additional details on this workflow, please consult the original Nipype tutorial (Notter 2020).

**Figure 5.**
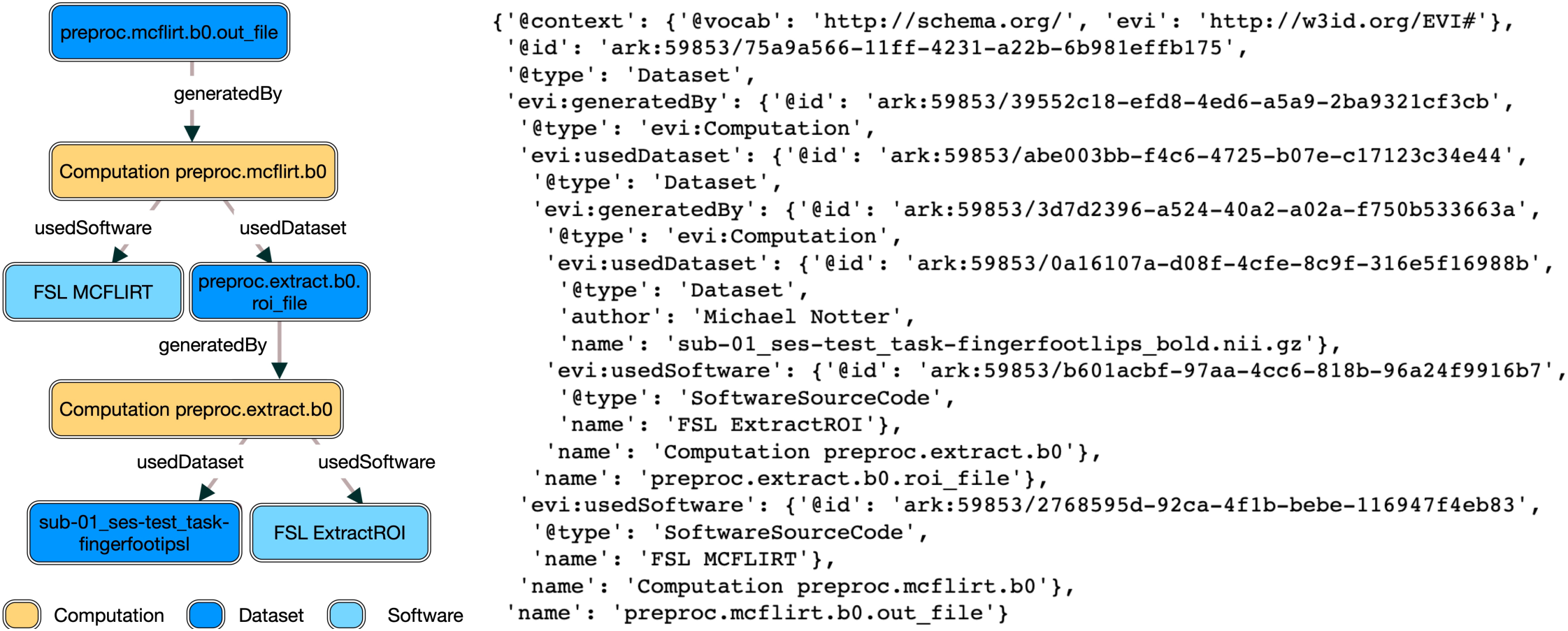
Evidence Graph visualization for the neuroimaging workflow execution.

## Discussion

FAIRSCAPE enables rapid construction of a shared digital commons environment and supports FAIRness within and outside that environment. It supports every requirement defined in the FAIR Principles at a detailed level, as defined in (Wilkinson et al. 2016), including a deep and comprehensive provenance model via Evidence Graphs, contributing to more transparent science and improved reusability of methods.

Scientific rigor depends on the transparency of methods (including software) and materials (including data). The historian of science Steven Shapin, described the approach developed with the first scientific journals as “virtual witnessing” (Shapin 1984), and this is still valid today. The typical scientific reader does not actually reproduce the experiment but is invited to review mentally every detail of how it was done to the extent that s/he becomes a “virtual witness” to an envisioned live demonstration. That is clearly how most people read scientific papers - except perhaps when they are citing them, in which case less care is often taken. Scientists are not really incentivized to replicate experiments; their discipline rewards novelty.

The ultimate validation of any claim once it has been accepted as reasonable on its face comes with support from multiple distinct angles, by different investigators; successful re-use of the materials and methods upon which it is based; and consistency with some body of theory. If the materials and methods are sufficiently transparent and thoroughly disclosed as to be reusable, and they cannot be made to work, or give bad results, that debunks the original experiments - precisely the way in which the promising-sounding STAP phenomenon was discredited (“RETRACTED ARTICLE: Stimulus-triggered fate conversion of somatic cells into pluripotency” 2014; Shiu 2014), before the elaborate formal effort of Riken to replicate the experiments (Ishii et al. 2014; RIKEN 2014).

As a first step then, it is not only a matter of reproducing experiments but also of producing transparent evidence that the experiments have been done correctly. This permits challenges to the procedures to develop over time, especially through re-use of materials (including data) and methods - which today significantly include software and computing environments. We definitely view these methods as being extensible to materials such as reagents, using the RRID approach; and to other computational disciplines.

## Conclusion

FAIRSCAPE is a reusable framework for scientific computations that provides a simplified interface for research users to an array of modern, dynamically scalable, cloud-based componentry. Our goal in developing FAIRSCAPE was to provide an ease-of-use (and re-use) incentive for researchers, while rendering all the artifacts marshalled to produce a result, and the evidence supporting them, Findable, Accessible, Interoperable, and Reusable. FAIRSCAPE can be used to construct, as we have done, a provenance-aware computational data lake or Commons. It supports transparent disclosure of the Evidence Graphs of computed results, with access to the persistent identifiers of the cited data or software, and to their stored metadata.

End-users do not need to learn a new programming language to use services provided by FAIRSCAPE. They require no additional special expertise, other than basic familiarity with Python and the skillsets they already possess in statistics, computational biology, machine learning, or other data science techniques. FAIRSCAPE provides an environment that makes large-scale computational work easier and results FAIRer. FAIRSCAPE is itself reusable and we have taken pains to provide well-documented straightforward installation procedures.

All resources on FAIRSCAPE are assigned identifiers which allow them to be shared. FAIRCAPE allows users to capture the complete evidence graph of the tasks performed. These evidence graphs show all steps of computations performed and the software and data that went into each computation. Evidence graphs, along with FAIRSCAPE’s other services, allow users to review and reproduce an experiment with significantly less overhead than other standard approaches. Users can see all computations that were performed, review the source code, and download all the data. This allows another party to reproduce the exact computations performed, apply the experimenter’s software to their own data, or apply their own methods to the experimenter’s data.

The optimal use case for a FAIRSCAPE installation is a local or multi-institution digital commons, in a managed Kubernetes environment. It can also be installed on high-end laptops for testing and development purposes as needed. We are actively looking for collaborators wishing to use, adapt, and co-develop this software.

FAIRSCAPE is not a tool for individual use, it is software for creating a high-efficiency collective environment. As a framework for sharing results and methods in the present, it also provides reliable deep provenance records across multiple computations, to support future reuse and to improve guarantees of reliability. This is the kind of effort we hope that centers and institutions will be increasingly making, to support reliable and shareable computational research results. We have seen increasing examples of philanthropic RFAs directing investment to this area. In our own research, we have found FAIRSCAPE’s methods invaluable in supporting highly productive computational research.

The major barriers to widespread adoption of digital commons environments, in our view, have been the relative non-reusability (despite claims to the contrary) of existing commons frameworks, and the additional effort required to manage a FAIR digital commons. We feel that FAIRSCAPE is a contribution to resolving the reusability issue, and may also help to simplify some digital commons management issues. Time will tell if the philanthropic investments in these areas noted above are continued. We hope they are.

We plan several enhancements in future research and development with this project, such as integration of additional workflow engines, including engines for genomic analysis. We intend to provide support for DataCite DOI and Software Heritage Identifier (SWHID) (Software Heritage Foundation 2020) registration; with metadata and data transfer to Dataverse instances, in future. Transfer of data, software, and metadata to long-term digital archives such as these, which are managed at the scale of universities or countries, is important in providing long-term access guarantees, beyond the life of an institutional center or institute.

Many projects involving overlapping groups have worked to address parts of the scale, accessibility, verifiability, reproducibility, and reuse problems targeted by FAIRSCAPE. Such challenges are in large part outcomes of the transition of biomedical and other scientific research from print to digital, and our increasing ability to generate data and to run computations on it at enormous scale. We make use of many of these prior approaches in our FAIRSCAPE framework, providing an integrated model for FAIRness and reproducibility, with ease of use incentives. We believe it will be a helpful tool for constructing provenance-aware FAIR digital commons, as part of an interoperating model for reproducible biomedical science.

## Information sharing statement

- Code for the microservices and the python client described in this paper are publicly available in the Zenodo repository at https://doi.org/10.5281/zenodo.4711204 and on GitHub at https://github.com/fairscape/fairscape. All versions are available under MIT license.
- Installation instructions, python Demo Notebooks, and API documentation are available at https://fairscape.github.io, also under MIT license.
- The MongoDB noSQL DB Community Version is available under MongoDB’s license terms at https://www.mongodb.com/try/download/community.
- The Stardog knowledge graph DB is available under Stardog’s license terms at https://www.stardog.com.
- The EVI ontology OWL2 vocabulary is available at https://w3id.org/EVI# under MIT license.

## Acknowledgements

We thank Chris Baker (University of New Brunswick), Caleb Crane (Mitsubishi Corporation), Mercè Crosas (Harvard University), Satra Ghosh (MIT), Carole Goble (University of Manchester), John Kunze (California Digital Library), Sherry Lake (University of Virginia), Maryann Martone (University of California San Diego), and Neal Magee (University of Virginia), for helpful discussions; and Neal Magee for technical assistance with the University of Virginia computing infrastructure. This work was supported in part by the U.S. National Institutes of Health, grants NIH OT3 OD025456-01, NIH 1U01HG009452, NIH R01-HD072071-05, and NIH U01-HL133708-01; and by a grant from the Coulter Foundation.

## Author Information

### Corresponding Author

correspondence to Timothy Clark

## Additional Information

- Data used in preparing this article was obtained from the University of Virginia Center for Advanced Medical Analytics and from OpenNeuro.org

## Rights and Permissions

This article is licensed under a Creative Commons Attribution 4.0 International License, which permits use, sharing, adaptation, distribution and reproduction in any medium or format, as long as you give appropriate credit to the original author(s) and the source, provide a link to the Creative Commons license, and indicate if changes were made. The images or other third-party material in this article are included in the article’s Creative Commons license, unless indicated otherwise in a credit line to the material. If material is not included in the article’s Creative Commons license and your intended use is not permitted by statutory regulation or exceeds the permitted use, you will need to obtain permission directly from the copyright holder. To view a copy of this license, visit http://creativecommons.org/licenses/by/4.0/.

